# TGF-β Signalling Drives Chemotaxis of Human Induced Pluripotent Stem Cell-Derived Cardiomyocytes in Response to MI Stimulus

**DOI:** 10.1101/2025.06.06.658033

**Authors:** Laura Deelen, Kazuya Kobayashi, Alia Hafiz Abbas Gasim, Fiona Lewis-McDougall, Ken Suzuki

**Author notes:** Joint corresponding authors. This study was funded by the British Heart Foundation (MRes/PhD DTP Programme, FS/19/62/34901).

## Abstract

Human induced pluripotent stem cell-derived cardiomyocytes (hiPSC-CMs) hold significant promise for cardiac regeneration therapies. However, the efficacy of such treatments depends on the ability of transplanted cells to migrate and integrate into the damaged myocardium, a process that remains poorly understood. In this study, we investigated the migratory behaviour of hiPSC-CMs using homogenised rat MI tissue to simulate myocardial infarction (MI) *in vitro*. Transwell migration assays demonstrated a concentration-dependent chemotactic response, with hiPSC-CM migration increasing up to threefold toward MI tissue homogenate. Wound healing assays further confirmed enhanced migration under MI-mimetic conditions. Bulk RNA sequencing revealed activation of the TGF-β signalling pathway as a key regulator of this response. Inhibition of TGF-β signalling, both pharmacologically and through antibody neutralisation, significantly reduced hiPSC-CM migration. These findings uncover a previously underappreciated chemotactic capability of hiPSC-CMs and identify TGF-β signalling as a central mediator, offering new mechanistic insights and potential therapeutic targets to improve the integration and efficacy of hiPSC-CM-based cardiac regeneration strategies.

## INTRODUCTION

Myocardial infarction (MI) is one of the most common causes of ischaemic heart failure (HF), for which the prognosis is only 50% within 5 years of diagnosis^1^. During MI, persistent coronary artery obstruction leads to the irreversible loss of billions of cardiac cells including cardiomyocytes (CMs). Currently the only established treatment for end-stage HF patients is heart transplantation or the permanent implantation of a left ventricular assist device as an emerging alternative^2,3^. However, these approaches have several limitations, including high costs, complications related to immunosuppression or anticoagulants, and psychological strain for patients. Myocardial regeneration therapy has drawn major attention over the last few decades as an innovative treatment to restore functional CMs after MI and improve prognosis through enhanced cardiac function. Human induced pluripotent stem cells (hiPSCs) have gained popularity as donor cells for myocardial regeneration therapy due to their unlimited self-renewal, autologous clinical potential and robust cardiomyogenic differentiation potency. Therefore, the transplantation of hiPSC-derived CMs (hiPSC-CMs) has become an active field of research for the treatment of post-MI HF.

Promising methods for delivery of replacement CMs to the heart include epicardial placement and intramyocardial injection of cell suspension or spheroids^4–6^. The migration of hiPSC-CMs following transplantation is a critical determinant of the success of cell-based cardiac regenerative therapies. Following epicardial delivery, a significant proportion of hiPSC-CMs have been shown to remain localized on the heart surface, failing to penetrate and integrate with the host myocardium ^7,8^. While intramyocardial injection offers more precise delivery into damaged myocardial regions, even this method relies on the transplanted cells’ ability to migrate within the myocardium to establish functional integration with surviving host cardiomyocytes. Such migration is essential for structural continuity, electrical coupling, and functional contribution to the regenerating heart. Therefore, limited intramyocardial migration remains a major barrier to the success of cell-based therapies.

Previous studies have begun to shed light on the migratory capacity of human stem cell-derived cardiomyocytes. Moyes et al. demonstrated that human embryonic stem cell-derived cardiomyocytes (hESC-CMs) exhibit enhanced migration *in vitro* in response to Wnt5a and fibronectin^9^. Another study identified BANCR, a primate-specific long non-coding RNA, as a positive regulator of hiPSC-CM migration^10^. Additional research on embryonic and neonatal CM migration has revealed that non-coding RNAs and developmentally regulated proteins may influence CM motility^11–14^.

Among the key pathways implicated in cardiomyocyte migration during regeneration is the TGF-β signalling pathway. In zebrafish, which possess innate cardiac regenerative capacity, SMAD3-dependent TGF-β signalling is essential for CM migration into damaged tissue and chemical inhibition of TGF-β signalling significantly impairs this migratory response^15^. Moreover, suppression of TGF-β–Smad3 signalling has been shown to inhibit an EMT-like response in CMs during zebrafish heart regeneration following ventricular ablation^16^.

It is known that MI triggers the release of numerous signalling molecules into the surrounding tissue, some of which may serve as chemoattractants for migratory cells. While adult CMs likely lack the capacity to respond to these cues, hiPSC-CMs due to their fetal-like phenotype, may retain an intrinsic ability to migrate in response to such signals, similar to regenerative responses observed in zebrafish^17^.

In this study, we aimed to investigate the migratory behaviour of hiPSC-CMs in response to infarct-associated signals. To this end, we employed transwell migration and wound healing assays to assess hiPSC-CM migration *in vitro*, using rat myocardial infarction MI tissue homogenate to mimic the post-MI microenvironment. Furthermore, we conducted bulk RNA sequencing on migrated and non-migrated hiPSC-CMs to identify key molecular pathways involved in this response.

Together, this study provides novel mechanistic insights into the chemotactic and migratory behaviour of hiPSC-CMs in the context of MI. These findings provide new insights into the migratory behaviour of hiPSC-CMs in response to infarct-associated cues and highlight molecular targets, such as TGF-β signalling, that could be modulated to enhance cell integration and improve functional outcomes in post-MI cardiac regenerative therapies.

## RESULTS

### HiPSC-CMs exhibit a concentration-dependent chemotactic response to rat MI tissue homogenate

Four hiPSC lines (hiPSC-A, B, C and D) were used in this study, each were confirmed to express surface and intracellular pluripotency markers and were karyotypically normal (Fig. S1). To generate hiPSC-CMs, we utilised the well-established protocol published by Burridge et. al^18^ or a commercial CM Differentiation Kit (STEMCELL Technologies) as an alternative. Spontaneous cell contractions were observed between days 7 and 11 post-induction of differentiation (Vid. S1).

Cardiomyocyte identity and differentiation efficiency were validated using flow cytometry, RT-PCR, and immunofluorescence staining. The Burridge protocol consistently yielded cTnT-positive cells across all four hiPSC lines (Fig. S2A), with protein expression of cTnT, NKX2.5, and α-actinin confirmed at day 15 (Fig. S2B). Gene expression profiling showed downregulation of pluripotency markers (POU5F1, NANOG, SOX2), transient upregulation of early mesoderm markers (MESP1, KIT), and increased expression of cardiomyocyte markers (NKX2-5, TNNT2, MYL7, MYL2) over time (Fig. S2C–G). For all downstream experiments, only differentiations achieving >60% cTnT+ cells by flow cytometry were included.

To assess whether MI tissue exerts a chemotactic effect on hiPSC-CMs, we developed a transwell migration (TWM) assay (Fig. 1A). Human iPSC-CM-A cells were seeded in the upper chamber and exposed to various dilutions of MI homogenate, derived from rat left ventricular tissue harvested 1-day post-MI. After 24 hours, migrated cells on the lower membrane surface were stained with DAPI and quantified (Fig. 1B–D). A significant, concentration-dependent increase in hiPSC-CM-A migration was observed in response to MI homogenate compared to control medium, with a peak 3-fold increase at a 1:10 dilution (Fig. 1C). Similarly, hiPSC-CM-B showed a robust 3-fold increase at the same dilution (Fig. 1D), and comparable results were seen in hiPSC-CM-C (Fig. S3).

**FIG 1.**
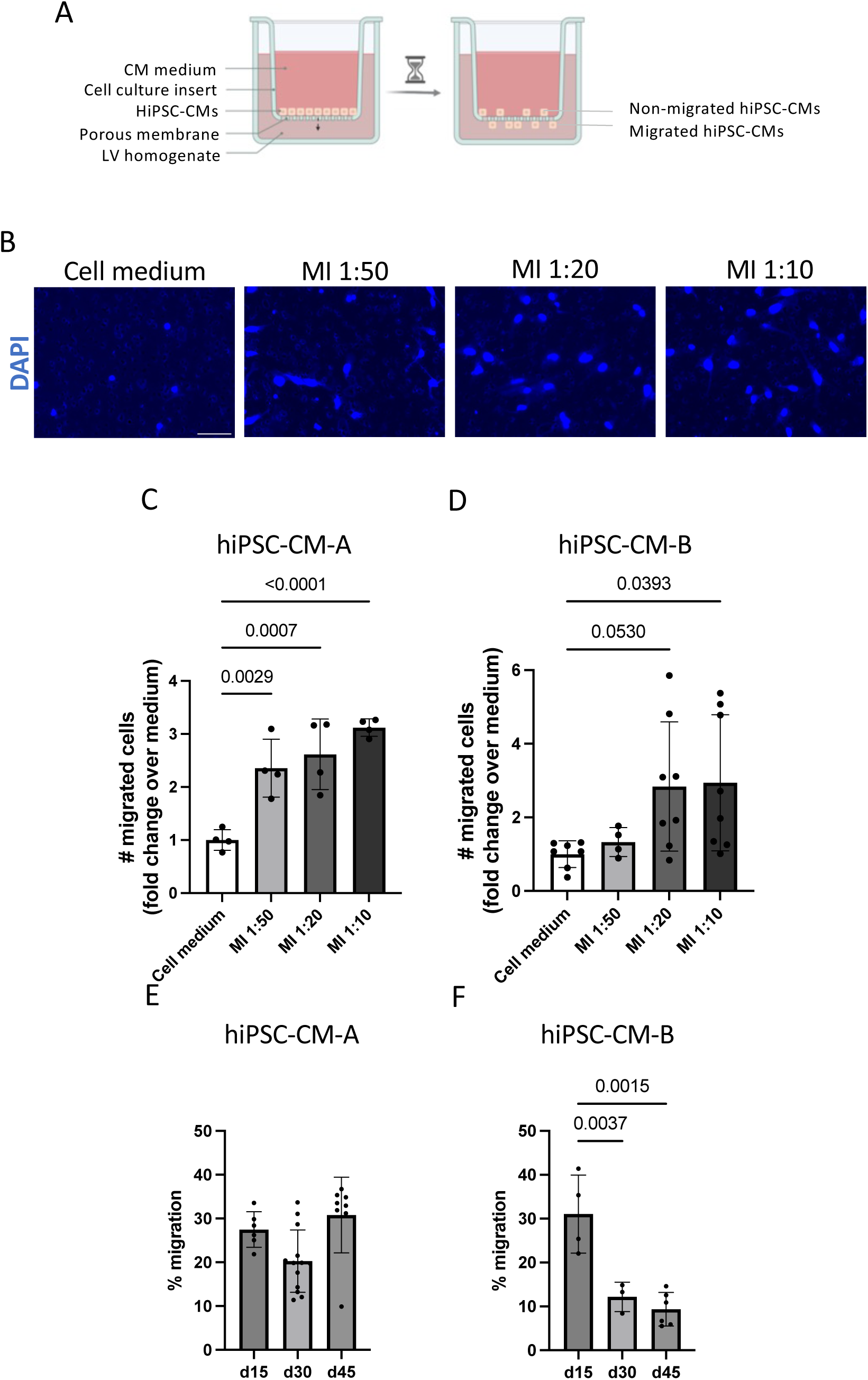
Human iPSC-CMs show a concentration-dependent chemotactic response to an MI stimulus that varies with maturation stage. A) Schematic of the transwell migration assay. B-D) HiPSC-CMs were seeded at day 15 and fixed and stained at 24h after incubation with cell medium, non-MI or MI homogenate at different dilutions. Non-migrated hiPSC-CMs on the top of the membrane were swabbed away and migrated hiPSC-CMs were imaged. B-C) hiPSC-CM-A, n=4. DAPI staining was used to visualise nuclei. Scale bar: 50 μm. D) hiPSC-CM-B, n=4-8, N=2. E-F) hiPSC-CM-A and hiPSC-CM-B were seeded at different days of differentiation and fixed and stained at 24h after incubation with MI homogenate (1:10), n= 6-10, N=1-3 (E), n=3-6, N=1-2 (F). Technical replicates are indicated with ‘n’ and biological replication with ‘N’. Statistical analyses were performed using one-way ANOVA with Dunnett’s correction. Error bars indicate mean ± SD.

We next explored whether the duration of hiPSC-CM culture influences their migratory capacity, given the known association between cardiomyocyte maturation and reduced motility^19,20^. Cells were assessed at various differentiation time points (day 15 to day 45) in the TWM assay. For hiPSC-CM-A, migration remained stable (~20–30%) across this time frame, with no statistically significant changes (Fig. 1E). In contrast, hiPSC-CM-B displayed a time-dependent decline in migration, decreasing from 28% at day 15 to just 9% by day 45 (Fig. 1F). These findings suggest cell line-specific differences in how prolonged culture impacts migratory ability.

Our transwell migration assays demonstrate that rat MI tissue homogenate induces a strong, concentration-dependent chemotactic response in hiPSC-CMs across multiple cell lines. Furthermore, cell line-specific and temporal factors influence the persistence of this migratory capacity during extended in vitro culture. These findings underscore the importance of accounting for both biological variability and maturation state when considering hiPSC-CMs for regenerative applications.

### An MI Stimulus Accelerates Collective hiPSC-CM Migration

To further investigate the impact of MI-associated cues on hiPSC-CM migration, we employed a wound healing assay to assess collective migration and wound closure dynamics. Human iPSC-CM-D monolayers were incubated with LV tissue homogenate from either MI or non-MI (sham) rats, and wound closure was assessed 4 hours after scratch injury (Fig. 2A–C).

**FIG 2.**
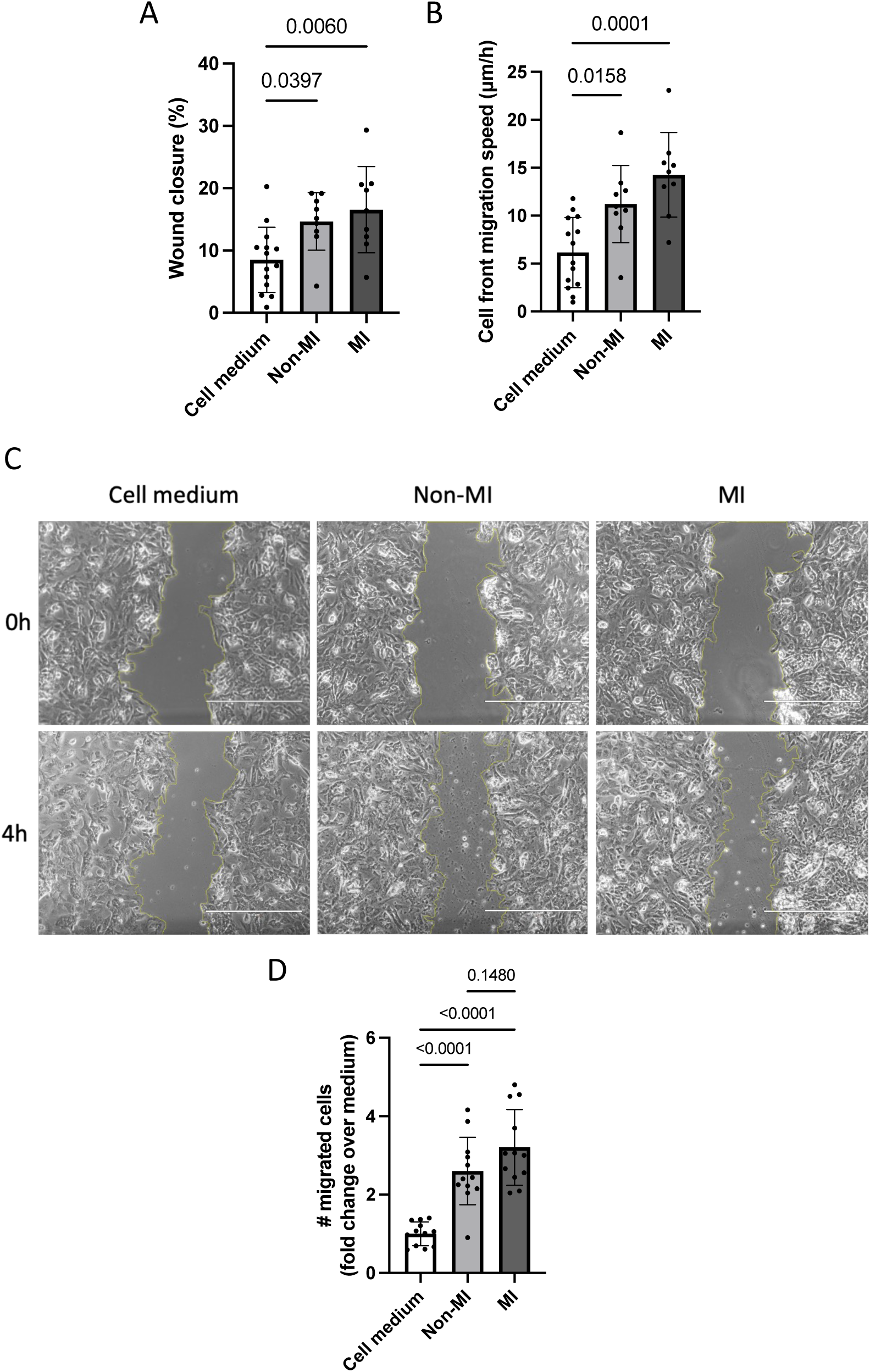
Human iPSC-CMs show a two-fold increase in migration when exposed to an MI environment. A-C) Day 15 hiPSC-CM-D (commercial kit) were replated to form a monolayer and a scratch was generated. HiPSC-CM-D were incubated with cell medium only (n=14) or supplemented with MI (1:10, n=9) or non-MI homogenate (1:10, n=9), wound closure (A) and cell front migration speed (B) were measured after 4h of incubation, N=2. C) Representative images of day 15 hiPSC-CM-D in the scratch wound assay. Scale bar: 1000 μm. D) Day 15 hiPSC-CM-A were seeded on transwell inserts and exposed to a 1:20 dilution of MI or non-MI homogenate for 4h (n=12). Technical replicates are indicated with ‘n’ and biological replication with ‘N’. Statistical analyses were performed using a one-way ANOVA followed by Tukey’s pairwise comparisons. Error bars indicate mean ± SD.

Exposure to MI homogenate (1:10 dilution) significantly enhanced wound closure compared to control medium (16.6 ± 6.9% vs. 8.5 ± 5.2%, respectively). This was accompanied by a substantial increase in the cell front migration speed, which more than doubled from 6.2 ± 3.7 μm/h in control conditions to 14.3 ± 4.4 μm/h in the presence of MI homogenate. Our observed migration speeds closely match those reported for hESC-CMs ^9^, reinforcing the biological validity of the model.

Interestingly, exposure to non-MI (sham) LV homogenate also resulted in enhanced wound closure (14.7 ± 4.6%) and increased migration speed (11.2 ± 4.0 μm/h) (Fig. 2A-C), though both metrics remained lower than those observed with MI homogenate. To further explore this observation, we conducted a transwell migration assay. Consistent with the wound healing results, non-MI homogenate induced a significant increase in hiPSC-CM migration relative to medium alone, but to a lesser extent than MI homogenate (Fig. 2D). This suggests that while sham homogenate does have chemokinetic effects, likely due to intracellular content release during tissue homogenisation, it lacks the full complement of injury-associated chemotactic cues present in the MI tissue.

### Migrated hiPSC-CMs exhibit a differential gene expression profile from non-migrated hiPSC-CMs

To identify pathways involved in hiPSC-CM migration, we employed a TWM assay in which day 15 hiPSC-CMs were exposed to rat MI homogenate for 24 hours. Migrated cells were collected from the lower surface of the membrane, while non-migrated cells were harvested from the upper surface. RNA was extracted from both populations for bulk RNA sequencing to comprehensively profile gene expression differences.

Principal component analysis (PCA) revealed clear segregation between the two hiPSC lines (hiPSC-CM-A and hiPSC-CM-B) along PC1, and between migrated and non-migrated samples of each line along PC2 (Fig. 3A). Together, PC1 and PC2 accounted for 91.53% of the total variance, indicating these components capture the main sources of gene expression variability. Notably, gene expression changes were more pronounced in hiPSC-CM-A than hiPSC-CM-B, highlighting inter-line variability.

**FIG 3.**
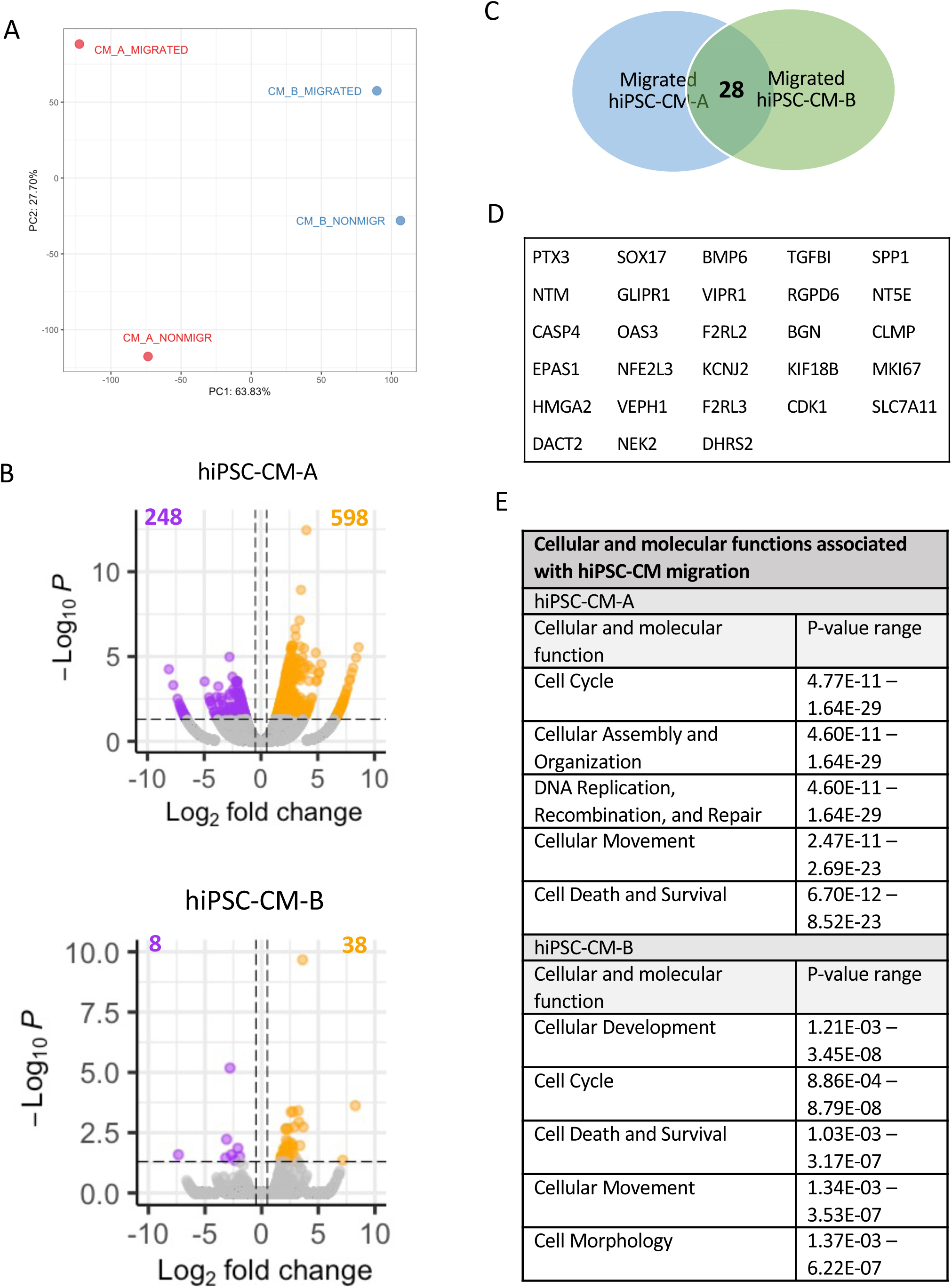
Migrated and non-migrated hiPSC-CMs represent distinct populations after 24h of incubation with MI homogenate. HiPSC-CM-A and hiPSC-CM-B were seeded on transwell inserts at day 15. After incubation overnight, the hiPSC-CMs were exposed to a 1:10 dilution of MI homogenate for 24h. Subsequently, migrated and non-migrated cells were harvested separately and analysed by bulk RNAseq. A) PCA plot. B) Differentially expressed genes in migrated cells versus non-migrated cells in hiPSC-CM-A and hiPSC-CM-B after 24h. Red dots indicate significantly upregulated genes, green dots indicate significantly downregulated genes and blue dots indicate genes which are not significantly differentially expressed. The cut-off parameters were chosen as log2FoldChange > |1| and padj or false discovery rate (FDR) <0.05. C-D) Combining the datasets resulted in a panel of 28 shared genes. E) Cellular and molecular functions associated with hiPSC-CM migration generated from the 24h dataset via Ingenuity Pathway Analysis.

To confirm cardiomyocyte identity, expression of canonical CM markers (TNNT2, NKX2-5, MYL7) was examined, showing comparable levels between migrated and non-migrated groups in both cell lines (Table S1). CM subtype markers indicated a heterogeneous population consisting of atrial (NR2F2), ventricular (MYL2, IRX4), and nodal (TBX18, HCN4) subtypes, consistent with expectations from the Burridge differentiation protocol (Fig. S2, Table S1). The immature phenotype of hiPSC-CMs was reflected by higher expression of fetal isoforms TNNI1 and MYH6 relative to adult isoforms TNNI3 and MYH7. Evaluation of potential contaminating cell types revealed low expression of cardiac fibroblast markers DDR2 and CD90 (THY1), and absence of endothelial marker CD31 in both lines (Table S1).

Following 24-hour incubation with MI homogenate, differential gene expression analysis identified 849 genes significantly altered in migrated versus non-migrated hiPSC-CM-A cells (598 upregulated and 248 downregulated; adjusted p-value < 0.05, log2 fold change > 1) (Fig. 3B). In contrast, only 46 genes were significantly differentially expressed in hiPSC-CM-B (38 upregulated, 8 downregulated). Integrating data from both lines yielded a consensus panel of 28 differentially expressed genes (DEGs) (Fig. 3C–D). Interestingly, proliferative markers MKI67 and CDK1 were present in this panel, suggesting that migrated hiPSC-CMs in both lines were more proliferative then non-migrating hiPSC-CMs.

From this panel, 14 genes of interest were selected for validation by RT-qPCR across hiPSC-CM-A, -B, and -C lines, confirming reproducible upregulation (Fig. S4). Gene Ontology (GO) enrichment analysis revealed that DEGs were predominantly associated with extracellular matrix organization, cell migration, and chemotaxis (Table S2). Ingenuity Pathway Analysis (IPA, QIAGEN) further identified cellular movement as a top molecular and cellular function linked to the differentially expressed genes in both datasets (Fig. 3F), supporting the ability of our model to uncover key regulators of hiPSC-CM migration

### Inhibition of TGF-β signalling reduces MI homogenate-induced hiPSC-CM migration

To investigate potential upstream regulators driving hiPSC-CM migration, we examined links between DEGs identified in our RNA-seq datasets. Among the 28 DEGs shared between migrated hiPSC-CM-A and hiPSC-CM-B, seven genes (SOX17, BMP6, TGFBI, SPP1, NT5E, HMGA2, and DACT2) have previously been directly associated with TGF-β signalling pathways ^23–28^. Furthermore, Ingenuity Pathway Analysis (IPA) identified TGF-β1 as a central upstream regulator associated with 12 of the 28 DEGs, implicating it in the observed migratory behaviour.

TGF-β signalling is known to be upregulated in the myocardium following MI ^29,30^ and plays a crucial role in heart regeneration in the zebrafish model, particularly by inducing an epithelial-to-mesenchymal transition (EMT)-like response in cardiomyocytes, which promotes migratory activity ^15, 16^. Notably, HMGA2, one of the DEGs identified in our study, is a well-characterized mediator of TGF-β-induced EMT ^28^.

To assess the functional relevance of TGF-β signalling in hiPSC-CM migration, we used the pharmacological inhibitor SB431542, which selectively blocks the kinase activity of the TGF-β type I receptor (ALK5) ^31^. In a scratch wound assay, treatment with SB431542 from 1 hour to 20 hours post-incubation with MI homogenate significantly reduced wound closure compared to MI homogenate alone (Fig. 4A). Similarly, cell front migration velocity was markedly decreased in the presence of SB431542, returning to baseline levels observed in control medium (Fig. 4B). Notably, the most pronounced effect occurred during the first 8 hours of incubation, suggesting that TGF-β signalling plays an early and critical role in initiating hiPSC-CM migration.

**FIG 4.**
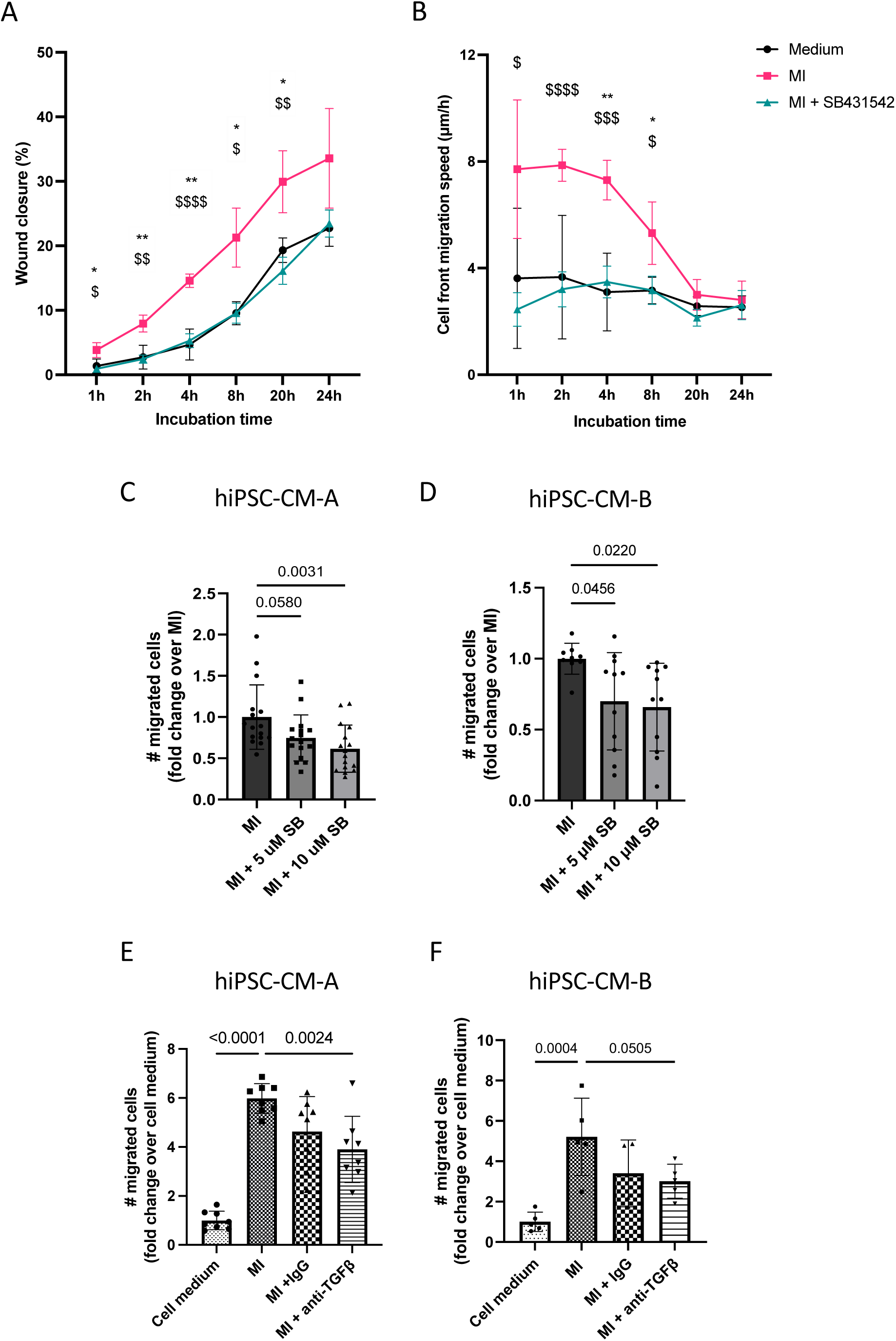
Inhibiting TGF-β signalling decreases MI-induced hiPSC-CM migration. A-B) Day 15 hiPSC-CM-A were seeded overnight and incubated with medium, MI 1:20 or MI 1:20 supplemented with 10 μM SB431542, n=4. *: P<0.05, **: P<0.01, MI vs. medium, $: P<0.05, $$: P<0.01, $$$: P<0.001, $$$$: P<0.0001, MI vs. MI + SB431542. C-D) HiPSC-CMs were seeded on transwell inserts and exposed to MI 1:20 only or with a 15-minute pre-incubation of 5 μM or 10 μM SB431542 prior to incubation with MI for 4h. C) hiPSC-CM-A, N=2 (d15 and d16), n=16. D) hiPSC-CM-B, N=3 (d15), n=10-11. E-F) D15 hiPSC-CMs were seeded on transwell inserts and exposed to cell medium, MI 1:20 only or MI with 20 ng/mL IgG or TGF-β neutralisation antibody for 4h. E) hiPSC-CM-A, n=12. F) hiPSC-CM-B, n=4-5. Technical replicates are indicated with ‘n’ and biological replication with ‘N’. Statistical analyses were performed using a one-way ANOVA followed by Tukey’s pairwise comparisons (C) or Dunnett’s correction (D-F). Error bars indicate mean ± SD.

We then confirmed the presence of TGF-β1 protein in the MI homogenate, measuring 1,014 ± 183 ng/mL, which was significantly higher than in non-MI (sham) homogenate (540 ± 146 ng/mL, Fig. S5). The detectable level of TGF-β1 in sham homogenate may account for its observed partial chemokinetic and chemotactic effects on hiPSC-CM migration (as described in Fig. 2).

To further confirm the involvement of TGF-β signalling, we performed a transwell migration assay. After 4 hours of incubation, the addition of 10 μM SB431542 to MI homogenate significantly reduced the number of migrated hiPSC-CMs (Fig. 4C–D). Similarly, the application of a neutralising anti–TGF-β1 antibody attenuated MI homogenate-induced migration. In hiPSC-CM-A, neutralisation led to a significant reduction in migrated cells (Fig. 4E), while hiPSC-CM-B showed a strong decreasing trend (p = 0.05, Fig. 4F).

These findings provide robust evidence that TGF-β1 is a key mediator of the migratory response of hiPSC-CMs to MI tissue stimuli. Pharmacological inhibition and antibody-based neutralisation both significantly impaired migration, supporting a causal role for TGF-β signalling in promoting hiPSC-CM motility. These results highlight TGF-β1 as a potential therapeutic target to modulate transplanted cell behaviour in cardiac regeneration strategies.

## DISCUSSION

This study provides, to our knowledge, the first comprehensive characterisation of the migratory capacity of hiPSC-CMs in response to factors derived from infarcted myocardium. Using two gold-standard *in vitro* migration models, transwell migration (TWM) and scratch wound assays, we demonstrated that hiPSC-CMs exhibit significant, concentration-dependent migratory responses when exposed to rat MI tissue homogenate. By employing multiple hiPSC lines, we confirmed the biological reproducibility of this migratory behaviour, establishing it as a previously unknown but relevant property of hiPSC-CMs in the context of MI.

The TWM model enabled the physical separation of migrated and non-migrated hiPSC-CM populations, allowing for bulk RNA sequencing to uncover gene expression differences potentially underpinning migration. Among the 28 DEGs common to two hiPSC lines, multiple were linked to the TGF-β signalling pathway, which was further validated through pharmacological and antibody-based inhibition. TGF-β has long been proposed as a chemoattractant ^32,33^ and is known to regulate migration in diverse cell types ^34–36^. Within the heart, it has been shown to influence cell migration during development and post-injury repair ^37^. In regenerative models such as zebrafish, TGF-β signalling plays a critical role in cardiac regeneration following injury, mediating cardiomyocyte migration through an EMT-like process ^15,16^. Our findings are consistent with these models and extend them by demonstrating that TGF-β1 also promotes hiPSC-CM migration in response to an MI stimulus.

We confirmed that MI homogenate contains significantly higher levels of TGF-β1 than non-MI (sham) homogenate and demonstrated that inhibiting TGF-β signalling with SB431542 or a TGF-β neutralising antibody significantly impaired hiPSC-CM migration. These results provide functional evidence that TGF-β1 is an active component of the infarcted myocardial microenvironment driving hiPSC-CM motility, and position it as a candidate target to enhance integration of transplanted cells into injured myocardium. However, the context-dependent nature of TGF-β signalling, which can elicit both beneficial and deleterious effects depending on cell type and injury phase, must be carefully considered ^38–41^.

Interestingly, our transcriptomic analysis also identified a proliferative gene expression profile in migrated hiPSC-CMs, including upregulation of MKI67 and CDK1. This suggests that migrated cells may possess enhanced proliferative potential, which could represent an added advantage following transplantation, contributing not only to cell engraftment but also to remuscularisation of infarcted tissue. Further studies are warranted to characterise and potentially harness this proliferative capacity *in vivo*.

We also examined the relationship between hiPSC-CM maturation and migration potential. As expected, prolonged culture led to either stable or reduced migratory capacity, depending on the cell line. While hiPSC-CM-A maintained consistent migration rates up to day 45 of differentiation, hiPSC-CM-B showed a marked decline over time. This suggests that cell line-intrinsic properties, alongside maturation status, influence migratory ability. Determining the optimal differentiation stage for transplantation will be critical, particularly as hiPSC-CMs undergo changes in metabolism, electrophysiology, and stress responses with maturation, all of which could impact therapeutic efficacy.

Our findings also raise important questions regarding cardiomyocyte subtype-specific migration. We observed early expression of the atrial marker MLC2a, which declined over time, while ventricular marker MLC2v increased, typical of CM maturation. However, there were no significant differences in the expression of MYL2 or MYL7 between migrated and non-migrated cells, suggesting that both atrial- and ventricular-like hiPSC-CMs (and double-positive cells) can respond to migratory cues. Nevertheless, recent protocols enabling the generation of highly enriched ventricular hiPSC-CMs ^42^ offer the opportunity to more precisely investigate the influence of CM subtype on migratory behaviour in future studies.

In summary, this study reveals novel insights into the migratory potential of hiPSC-CMs, identifies TGF-β1 as a key mediator of this response, and highlights the importance of cell-intrinsic properties, maturation state, and myocardial context in modulating migration. These findings contribute to our understanding of how transplanted hiPSC-CMs might interact with the post-MI environment, with potential implications for improving cell therapy strategies aimed at cardiac regeneration. Targeting the migratory behaviour of hiPSC-CMs will enable us to promote effective cell engraftment and integration, ultimately supporting functional repair of the injured heart.

## METHODS AND MATERIALS

### Human iPSC culture

Four hiPSC lines were obtained from the European Bank for Induced Pluripotent Stem Cells (EBiSC) and maintained in feeder-free, serum-free 2D culture at 37°C and 5% CO_2_. For CM differentiation, hiPSCs were grown in E8 Flex medium (Thermofisher) on recombinant human vitronectin (rhVTN)-coated plates for 3-4 days until reaching approximately ~85% confluency. Differentiation into cardiomyocytes was initiated using a protocol adapted from Burridge et al.¹⁸ Briefly, hiPSCs were incubated in CDM3 medium; RPMI 1640 supplemented with 75 mg/mL recombinant human albumin (Merck) and 64 mg/mL L-ascorbic acid 2-phosphate (Merck) with the addition of 6 μM CHIR99021 (Cambridge Bioscience) for 48 hours. On day 2, the medium was replaced with CDM3 containing 2 μM Wnt-C59, and from day 4 onwards cells were maintained in CDM3 alone, with media refreshed every two days. Alternatively, where indicated, hiPSC-CMs were generated using a commercially available Cardiomyocyte Differentiation Kit (STEMCELL Technologies), following the manufacturer’s instructions.

### Heart tissue homogenisation

Adult male Sprague-Dawley rats underwent either MI induction via left coronary artery (LCA) ligation or a sham surgical procedure. For MI, the LCA was ligated at the level of the lower edge of the left atrial appendage via left thoracotomy under isoflurane anaesthesia and mechanical ventilation. Successful induction of ischaemia was confirmed by regional epicardial colour change and the presence of dyskinesia. Following chest and skin closure, animals were extubated and allowed to recover in their cage. Sham-operated animals underwent an open-chest procedure without LCA ligation. All animals were sacrificed 24 hours post-surgery. The left ventricular (LV) wall was then harvested, either freshly homogenised or snap-frozen in liquid nitrogen and stored at –80 °C for later use.

Tissue homogenisation was performed PBS supplemented with protease inhibitors (cOmplete™, EDTA-free Protease Inhibitor Cocktail; Sigma-Aldrich) using a Precellys 24 tissue homogeniser. Homogenates were centrifuged at 6,000 × g for 15 minutes, and the supernatant was collected for immediate use or stored at –80 °C for subsequent experiments.

All animal procedures were approved by the institutional ethics committee at Queen Mary University of London and the UK Home Office. Experiments conformed to the *Principles of Laboratory Animal Care* (National Society for Medical Research) and the *Guide for the Care and Use of Laboratory Animals* (U.S. National Institutes of Health, 1996).

### Wound healing assay

Human iPSC-CMs were seeded at a density of ≥ 2.5 × 10^5^ cells/cm^2^ to form a confluent monolayer. To promote cell attachment, hiPSC-CMs were incubated overnight with 10 μM Y-27632 (Rock inhibitor; Cambridge BioScience). The following day, a scratch wound was generated across the hiPSC-CM monolayer using a sterile P1000 tip. After washing with PBS to remove detached cells, hiPSC-CMs were incubated for up to 24 hours with either cell media alone or medium supplemented with LV homogenate from MI or sham operated rat hearts (diluted as specified).

Cell migration was assessed by capturing images at defined time points using either a Keyence BZ-X810 microscope or an EVOS XL Core Cell Imaging System. Quantitative analysis was performed using ImageJ software. Percentage wound closure was calculated as follows;

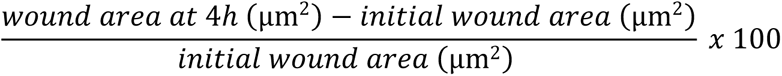

Cell front migration was calculated as follows;

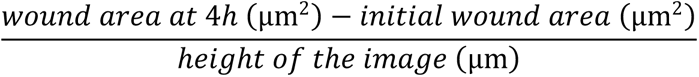

in which the height of the image represents the y-axis, to obtain the difference in horizontal distance (x-axis), i.e., the cell front migration (μm). This number is then divided by the incubation time to obtain the cell front migration speed (μm/h).

### Transwell migration assay

All TWM assays were performed in 24-well plates. Transwell inserts with 8 μM pore size (Starstedt) were pre-coated on both sides with ~22 μg/cm^2^ Geltrex Ready to Use (Thermofisher) for 1h at 37°C. After aspiration of the residual coating, 1 × 10^5^ hiPSC-CMs per insert were seeded into the upper chamber in cardiomyocyte support medium supplemented with 10 μM Y-27632 (Rock inhibitor)and incubated overnight to allow cell attachment. The next day, the medium in the lower chamber was replaced with cell medium alone or medium supplemented with LV homogenate from MI or sham operated rat hearts at the indicated dilution. After 24 hours of incubation, cells were fixed in 4% paraformaldehyde for 15 mins at room temperature (RT) followed by a PBS wash and incubation with DAPI for 15 mins at RT. To ensure that only migrated cells on the lower surface were visualised, the upper surface of the insert was gently swabbed with a cotton applicator to remove non-migrated cells. The membranes were then carefully excised and mounted onto glass microscope slides. Images were acquired on a Keyence BZ-X180 or Zeiss LSM800 microscope. Quantification of migrated cells was performed using ImageJ software.

### ELISA

TGF-β1 levels were measured in homogentes from sham and MI tissues using a TGF-beta 1 Quantikine ELISA kit (R&D, Cat #DB100C). To activate latent TGF-β1, tissue supernatents were incubated with 1M HCl for 10 mins, followed by neutralisation with 1.2M NaOH/0.5M HEPES. Activated samples or standards were added to wells pre-coated with TGF-β1 capture antibody, along with the assay diluent. After a 2h incubation at RT, plates werewashed four times before addition of the TGF-β1 conjugate. Plates were then incubated for a further 2 hours, followed by four washes. Substrate solution was then added and incubated for 30 mins at RT in the dark. The reaction was stopped by the addition of Stop Solution and absorbance was measured at with wavelength correction at 540 nm using a microplate reader.

### Bulk RNAseq

Human iPSCs were seeded onto transwell inserts at day 15 of CM differentiation, as previously described. After 24 hours of incubation with a 1:10 dilution of left ventricular homogenate from a MI rat heart in the lower compartment, culture media was aspirated and the inserts washed in PBS. To isolate migrating and non-migrating cells, the upper and lower surfaces of the inserts were swabbed separately.

.RNA was extracted by lysing the cells directly on the inserts using RLT lysis buffer from the Qiagen RNeasy mini kit, following the manufacturer’sinstructions. Samples with RNA integrity numbers (RIN)≥ 9.4 and a minimum input of 100ng of total RNA were used for bulk RNAseq (20 million reads per samples). Library preparation and paired-end 150 bp sequencing (PE150) were performed by Novogene Ltd, who also conducted downstream data analysis.

### Immunocytochemistry

Human iPSCs were harvested by incubation with EDTA for 3-5 mins, followed by gentle detachment with culture media. Cells were seeded onto 16 mm coverslips in 12 well plates pre-coated with rhVTN and cultured for 2-4 days before fixation with 4% paraformaldehyde in PBS for 15 min at 37°C.

Ahead of immunostaining, hiPSC-CMs were replated overnight using the CM dissociation kit (Thermofisher). Briefly, wells were washed twice with PBS before incubation with CMdissociation medium at 37°C for 10-15 mins. Detached cells were collected usinga 10 mL serological pipette and transferred into tubes containing CM support medium. After centrifugation at 300 g for 5 min, the supernatant was removed, and the cell pellet was resuspended in CM support medium supplemented with 5 µM Y-27632. Cells were seeded into a new rhVTN-coated plate at the desired density (≥ 2.5 × 10^5^ cells/cm^2^) and allowed to attach overnight. The following day, cells were fixed with 4% paraformaldehyde in PBS for 15 mins at 37°C.

After fixation, hiPSCs and hiPSC-CMs were washed and permeabilised with 0.1% Triton-X-100 inPBS for 10 min at RT, then blocked using 10% donkey serum/PBS for 1h at RT. Primary antibodies were applied overnight at 4°C in the dark. After washing, cells were incubated with appropriate secondary antibodies for 30 mins at 37°C. DAPI was used as a nuclear counterstain. All wash steps were performed with 0.1% Tween in PBS. Imaging was conducted using a Zeiss LSM 800 or Keyence BZ-X810 microscope. The antibodies used for immunostaining can be found in Supplementary Tables S4 and S5.

### Flow cytometry

Human iPSCs were harvested by incubation with EDTA for 3-5 mins followed by aspiration, followed by gentle detachment with culture media. Cells were then transferred to 15 mL tubes and centrifuged at 200 g for 5 mins. The supernatant was aspirated, and cells were resuspended into single cells in filtered FACS buffer (PBS, 4% w/v bovine serum albumin (BSA), 10% v/v Penicillin-Streptomycin (Pen-Strep), 4% v/v EDTA). After repeating the last step, the cells were stained for 30 mins at RT. The antibodies used can be found in Supplementary Table S4–5.

HiPSC-CMs were dissociated with the Cardiomyocyte Dissociation Kit (see above) and stained with the Inside Stain Kit (Miltenyi Biotech) according to the manufacturer’s protocol. Briefly, dissociated cells are counted with Trypan Blue and up to 1 million cells are washed by adding FACS buffer. After centrifugation at 200 g for 10 mins, the cells are resuspended in 1:1 FACS buffer/Inside Fix buffer. After 20 mins of incubation in the dark at RT, the cells are washed with FACS buffer and centrifuged at 200 g for 5 mins. The washing step is repeated in Inside Perm. The cells were directly stained in Inside Perm buffer for 10-30 mins at RT, or indirectly for 1h at RT and with secondary antibodies for 30 mins at RT in the dark. The antibodies used can be found in Supplementary Table S4–5.

HiPSCs/hiPSC-CMs were subsequently washed and resuspended in incubation medium and were kept on ice until analysis. Data was acquired on a BD LSRFortessa instrument and analysed using FlowJo software.

### Real-time quantitative PCR (RT-qPCR)

Total RNA isolation was performed using the RNeasy mini kit (Qiagen) according to the manufacturer’s protocol. Briefly, harvested cells or heart homogenates were lysed in RLT lysis buffer for 5 mins at RT. The resulting cell lysate was transferred to a QIAShredder column and was centrifuged at full speed for 3 mins. Next, 70% ethanol was added to the lysate, which was transferred to an RNeasy silica column, followed by centrifugation at 1.2 × 10^4^ rpm for 15s. This was followed by a washing step in RW1 wash buffer and a DNase I incubation for 15 mins at RT, followed by another wash step with RW1 and twice with RPE, followed by dry centrifugation of the column at 1.2 × 10^4^ rpm for 1 min. The RNA was eluted in RNase free ddH2O for 2 min at RT followed by centrifugation at 1.2 × 10^4^ rpm for 1 min. Eluted RNA was kept on ice until further use or stored at −80°C for future use. RNA was subsequently reverse transcribed into cDNA with the High Capacity Reverse Transcription Kit (Thermofisher, 4368814) with the following thermal cycler settings: 10 min at 25°C, 120 mins at 37°C, 5 mins at 85°C. Gene expression was quantified by real-time PCR of 10 ng cDNA of each sample with the PowerUp SYBR Green supermix (Thermofisher, A257420). Primers for each gene are given in Table S6. β-actin and GAPDH were used as housekeeping genes. Of note, the use of β-actin and GAPDH as reference genes has been critiqued as these genes show variable levels of expression in different tissues during development including the heart, which might compromise their role in gene expression normalisation (Lin & Redies, 2012).

The plate was transferred to a QuantStudio 7.0 Real-Time PCR machine (Applied Biosystems) and the following parameters were applied: 5 mins 95°C, 40 cycles of 15s at 95°C, 30s at 60°C and 30s at 72°C, followed by a melt curve analysis from 55-95°C.

### Statistics

All data was analysed using Graphpad Prism 9.0 software (Graphpad, San Diego, CA). Data is represented as mean ± standard deviation (SD). Statistical analysis was performed on data with a replicate number of 3 and above. Technical replicates are indicated with ‘n’ and biological replication with ‘N’, as detailed in the figure legends. For the comparison between two groups, two-tailed unpaired Student’s t-tests were performed. Multiple group comparisons were performed with one-way or two-way analysis of variance (ANOVA) with Tukey’s post hoc test correction (3 groups) or Dunnett’s correction (>3 groups). If the number of replicates was not equal between groups, a mixed-effects analysis was performed instead of an ANOVA, as the latter cannot deal with missing values. Statistical significance was accepted at p <0.05.

## AUTHOR CONTRIBUTIONS

KS, FLM and LD conceived and designed the study. LD performed the majority of the experiments. KK and AHAG contributed to the execution of select experiments. LD, KS and FLM analysed the data and wrote the manuscript. All the authors read and approved the manuscript.

---

**FIG S1:**
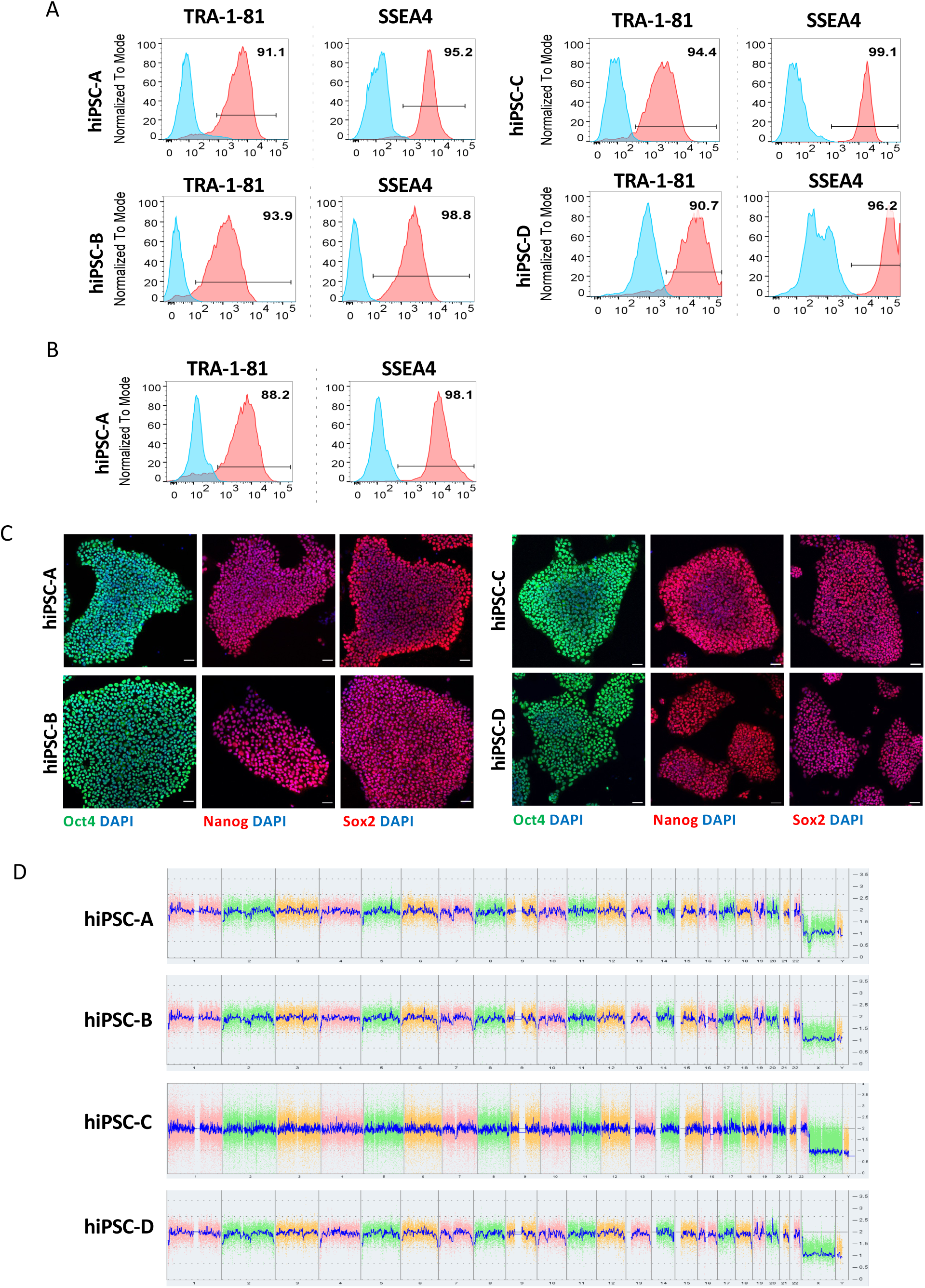
Pluripotency and karyotype normality of hiPSC lines. A) Flow cytometric analysis of pluripotency markers, TRA-1-81 and SSEA4 of hiPSC-A (P47), hiPSC-B (P34), hiPSC-C (P34) and hiPSC-D (P41). Samples are shown in red and isotype controls are shown in blue. B) Flow cytometric analysis of TRA-1-81 and SSEA4 of hiPSC-A at passage 77. C) Representative micrographs showing immunolabelling of pluripotency markers Oct-4 (green), Nanog (red) and Sox2 (red), nuclei are labelled with DAPI (blue). Scalebar: 50 µm. D) hiPSCs A-D were negative for karyotypic abnormalities confirmed by KaryoStat analysis at passage 54, 47, 36 and 44, respectively. The whole genome view displays all somatic and sex chromosomes in one frame with copy number levels. The pink, green and yellow colours indicate the raw signal for each individual chromosome probe, while the blue signal represents the normalized probe signal which is used to identify copy number and potential aberrations deviating from a normal copy number state (CN = 2).

**FIG S2:**
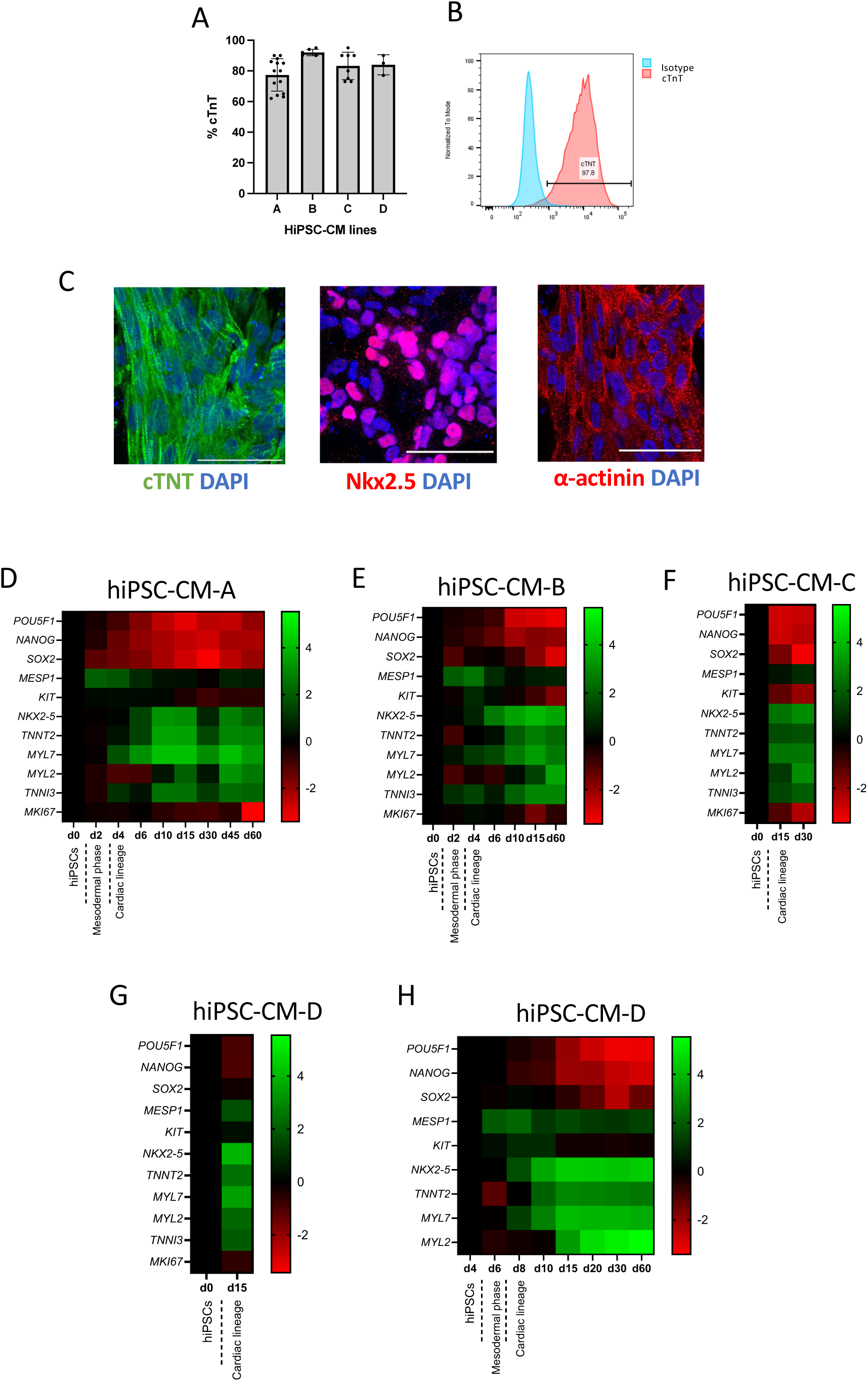
Expression of pluripotency and cardiac markers during hiPSC-CM differentiation using the Burridge protocol and commercial kit. A) Differentiation efficiency measured by cTnT expression in hiPSC-CM-A-D lines using the Burridge et al. protocol. B) Representative graph of cTnT expression measured in hiPSC-CM-B at day 15. C) Representative micrographs showing immunolabelling for cardiac markers at day 15 of hiPSC-CM-D differentiation using the commercial kit. Scalebar: 50 µm. D-H). Heat maps illustrating quantitative RT-qPCR analysis of pluripotent, mesodermal, and cardiac marker expression during hiPSC-CM differentiation of hiPSC-CM-A (D), hiPSC-CM-B (E), hiPSC-CM-C (F) and hiPSC-CM-D (G) using the Burridge protocol and hiPSC-CM-D using the commercial kit (H). Values are represented as log fold change and represent the average of 1-3 samples.

**Fig S3:**
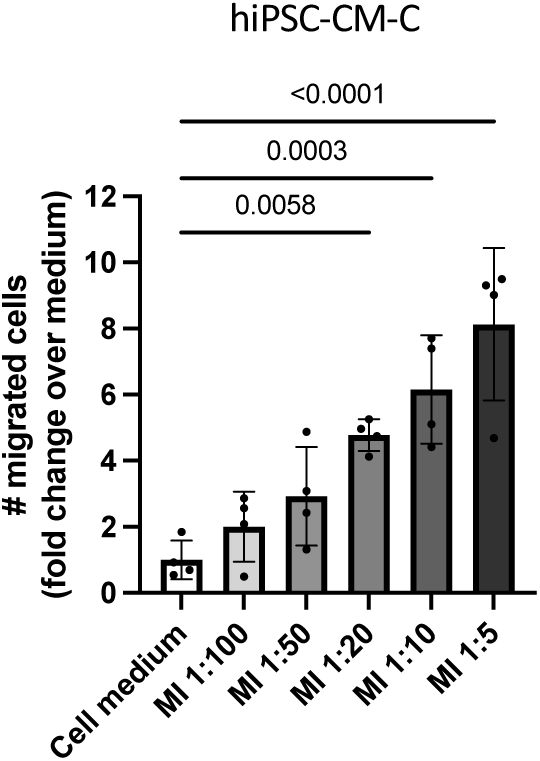
MI homogenate has a concentration-dependent effect on hiPSC-CM-C migration. HiPSC-CM-C were seeded onto transwells at day 15 and fixed and stained at 24h after incubation with cell medium, non-MI or MI homogenate at different dilutions. Non-migrated hiPSC-CMs on the top of the membrane were swabbed away and migrated hiPSC-CMs were imaged. n=4. Statistical analysis was performed using one-way ANOVA followed by Dunnett’s correction. Error bars indicate mean ± SD.

**FIG S4:**
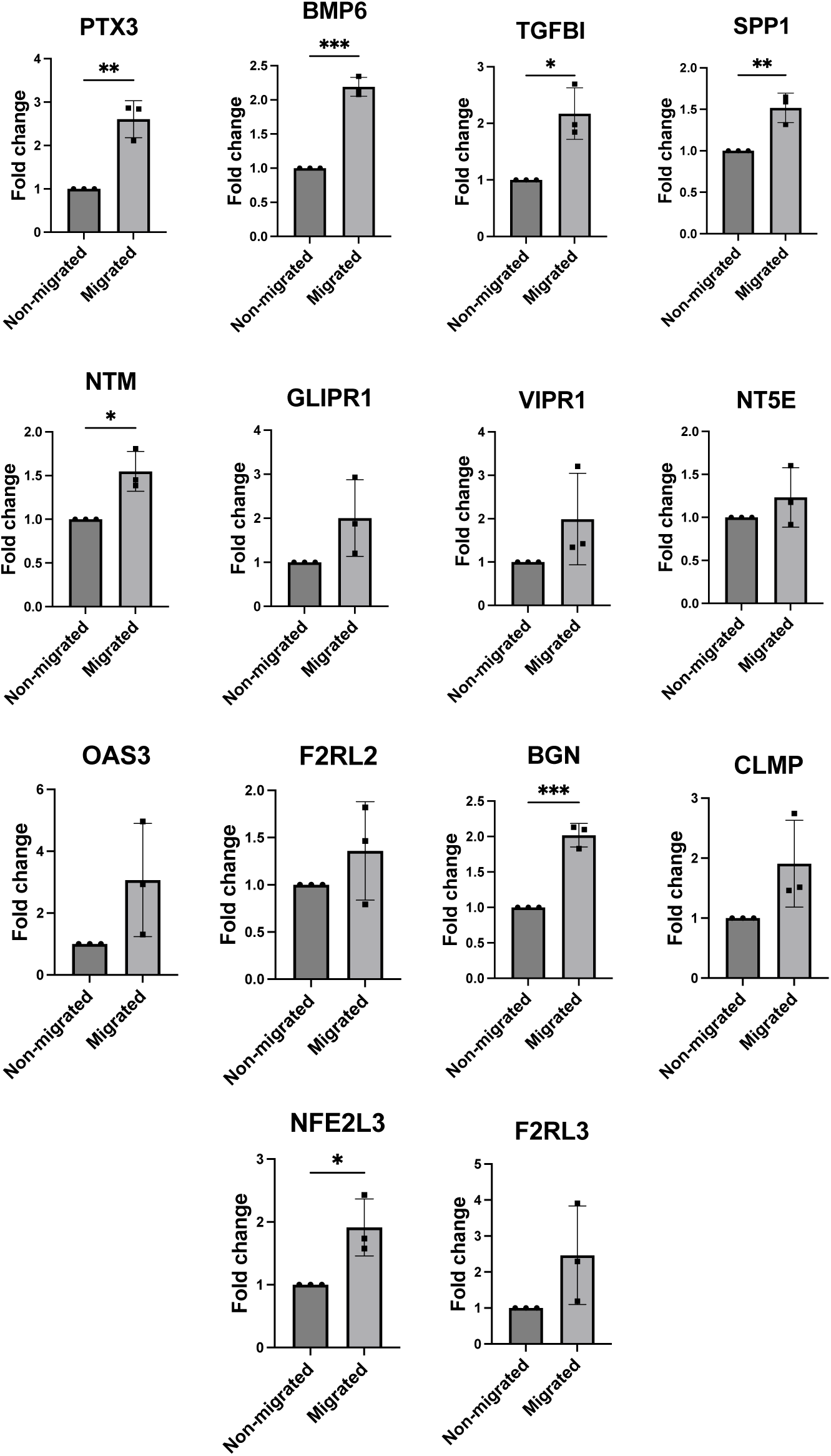
Differentially expressed gene verification comparing migrated hiPSC-CMs to their non-migrated counterparts. HiPSC-CM-A,-B and -C were seeded at day 15 on transwell inserts and exposed to MI 1:10 for 24h after which the non-migrated and migrated cells were harvested separately. RT-qPCR confirmed the upregulation of all 14 genes of interest in migrated cells compared to their non-migrated counterparts in all three biological replicates. Statistical analyses were performed using two-tailed, unpaired t-tests. Error bars indicate mean ± SD. *: P ≤ 0.05, **: P ≤ 0.01, ***: P≤ 0.001.

**FIG S5:**
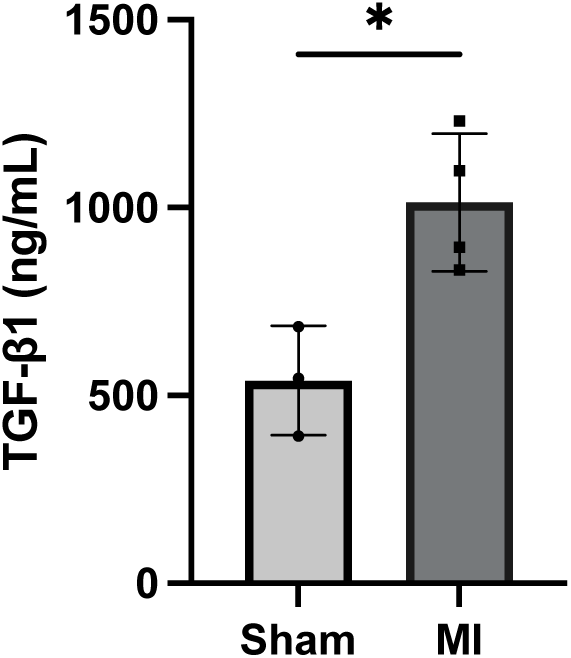
TGF-β1 levels in sham and MI rat left ventricular homogenates. TGF-β1 was measured in sham (n=3) or MI (n=4) LV homogenate samples by ELISA. Statistical analysis was performed using a two-tailed unpaired t-test. *, P < 0.05.

**Table S1:** Gene expression profiles of hiPSC-CMs analysed for bulk RNAseq after 24h of incubation with MI homogenate (1:10). Values represent normalised read counts. Padj = adjusted p-value or false discovery rate (FDR).

| Gene | hiPSC-CM-A |  |  | hiPSC-CM-B |  |  |
| --- | --- | --- | --- | --- | --- | --- |
|  | Migrated | Non-migrated | padj | Migrated | Non-migrated | padj |
| *TNNT2* | 8071 | 18864 | 0.07 | 11685 | 16025 | 1 |
| *NKX2-5* | 3044 | 4933 | 0.48 | 3892 | 4316 | 1 |
| *MYL7* | 18495 | 44382 | 0.06 | 26798 | 38134 | 1 |
| *NR2F2* | 1109 | 487 | 0.10 | 746 | 540 | 1 |
| *MYL2* | 60 | 142 | 0.14 | 91 | 123 | 1 |
| *IRX4* | 126 | 254 | 0.23 | 291 | 257 | 1 |
| *TBX18* | 1193 | 779 | 0.56 | 206 | 85 | 0.59 |
| *HCN4* | 1458 | 2882 | 0.21 | 1097 | 1643 | 1 |
| *TNNI3* | 231 | 687 | 0.01 | 280 | 515 | 1 |
| *TNNI1* | 2620 | 6508 | 0.05 | 3499 | 5198 | 1 |
| *MYH6* | 26392 | 88865 | 0.003 | 14032 | 32149 | 0.53 |
| *MYH7* | 4489 | 16524 | 0.001 | 2285 | 4259 | 1 |
| *DDR2* | 806 | 403 | 0.21 | 599 | 284 | 0.84 |
| *THY1* | 368 | 149 | 0.07 | 30 | 5 | 0.23 |
| *CD31* | - | - |  | - | - |  |

**Table S2:**
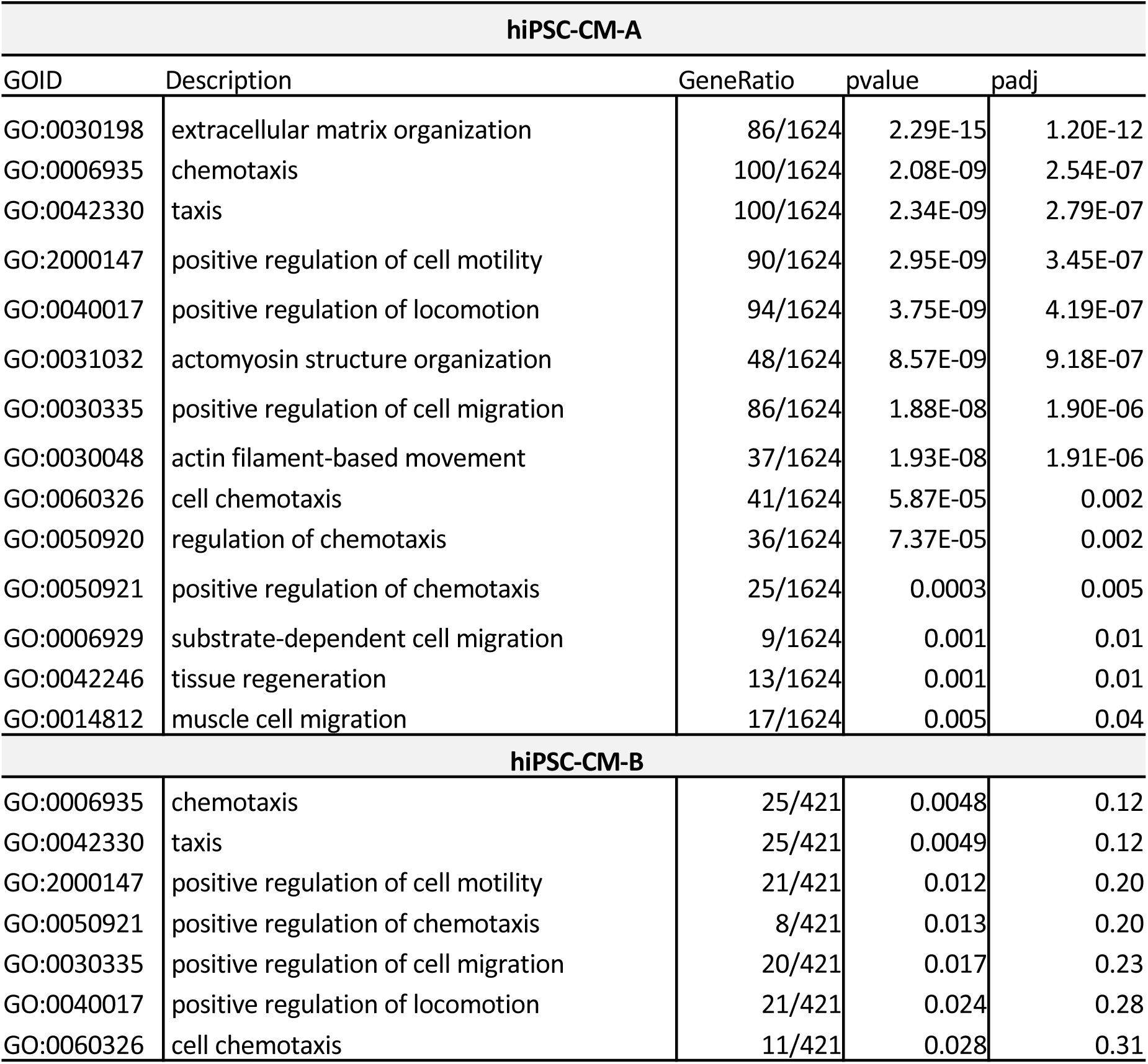
Gene Ontology (GO) analysis of migrated versus non-migrated hiPSC-CM-Bs and hiPSC-CM-Cs after 24h of incubation with MI homogenate.

**Table S3:**
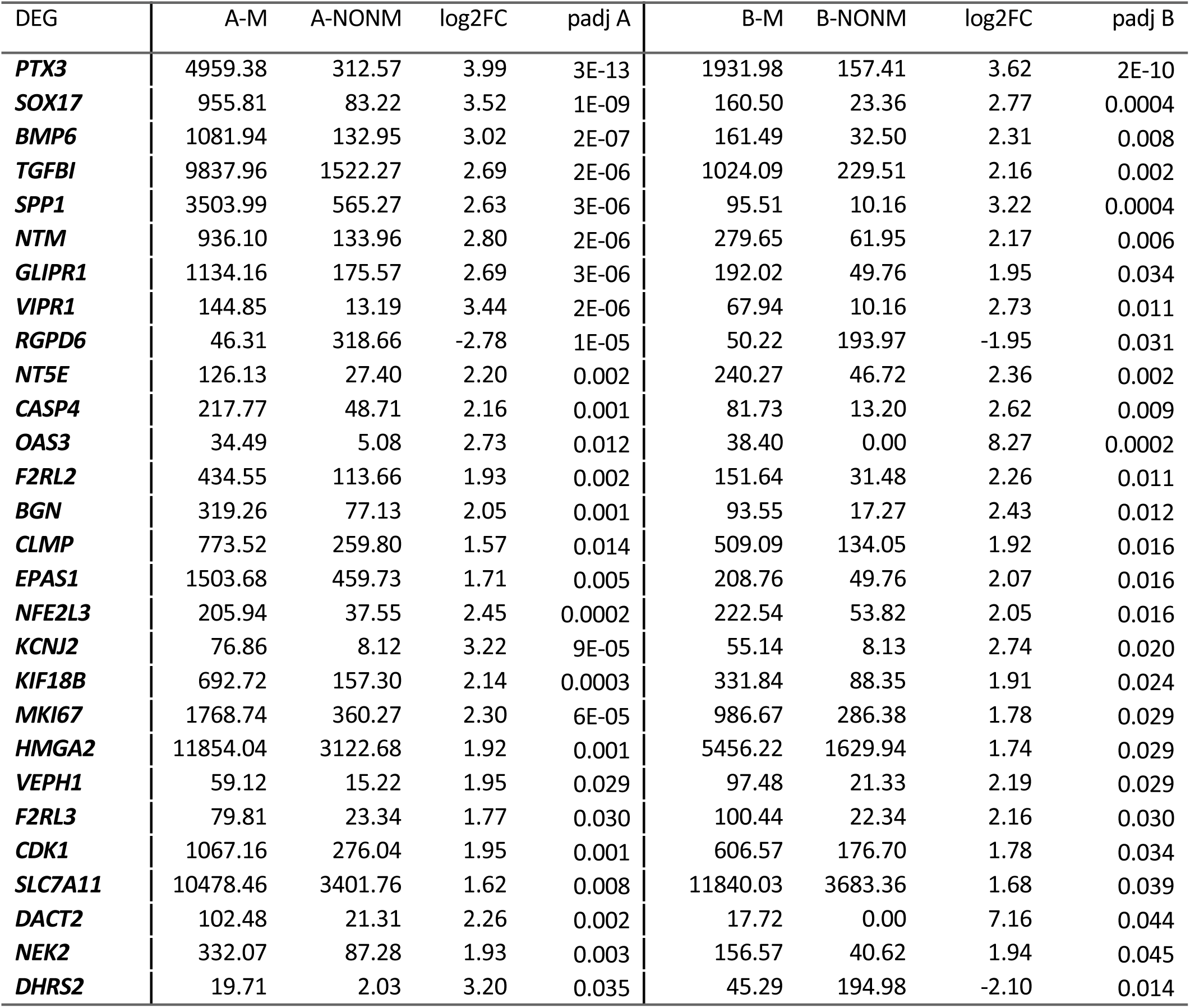
Differentially expressed genes (DEGs) shared between migrated (A-M,B-M) versus non-migrated hiPSC-CM-A (A-NONM) and hiPSC-CM-B (B-NONM) after 24h of incubation with MI homogenate diluted 1:10. Values represent as normalised read counts. Padj = adjusted p-value or false discovery rate (FDR).

**Table S4.** List of unconjugated antibodies used in this study. All antibodies are reactive against human. ICC: immunohistochemistry, WB: western blot, FC: flow cytometry.

|  |  |  |  |  |
| --- | --- | --- | --- | --- |
| Target | Host species | Source | Application | Working concentration |
| Oct3/4 | Goat | R&D, #AF1759 | ICC | 1:50 |
| Nanog | Goat | R&D, #AF1997 | ICC | 1:50 |
| Sox2 | Rabbit | Abcam, #Ab97959 | ICC | 1:500 |
| Nkx2.5 | Goat | R&D, #AF2444 | ICC | 1:20 |
| cTNT | Mouse | Thermofisher, #MA5-12960 | ICC, WB, FC | 1:50 |
| Sarcomeric alpha-actinin | Mouse | Sigma (Merck), #A7811 | ICC | 1:250 |
| MLC2v | Rabbit | Proteintech, #10906-1-AP | FC | 1:400 |

**Table S5.** List of conjugated antibodies used in this study. ICC: immuno-histochemistry, WB: western blot, FC: flow cytometry.

|  |  |  |  |  |
| --- | --- | --- | --- | --- |
| Target | Conjugate | Source | Application | Working concentration |
| Anti-mouse | AF488 | Thermofisher, #A21202 | ICC | 1:500 |
| Anti-mouse | AF647 | Thermofisher, #A21242 | FC | 1:500 |
| Anti-goat | AF488 | Thermofisher, #A11055 | ICC | 1:500 |
| Anti-goat | AF594 | Thermofisher, #A11058 | ICC | 1:500 |
| Anti-rabbit | AF488 | Thermofisher, #A21206 | ICC | 1:500 |
| Anti-rabbit | AF555 | Thermofisher, #A32732 | FC | 1:500 |
| Anti-rabbit | AF594 | Thermofisher, #A11037 | ICC | 1:500 |
| Anti-human TRA-1-81 (IgM, κ) | AF488 | Biolegend, #330710 | FC | 1:10 |
| Anti-human isotype (IgM, κ) | AF488 | Biolegend, #401617 | FC | 1:10 |
| Anti-human SSEA4 (IgG3) | AF488 | Thermofisher, #53-8843-42 | FC | 1:10 |
| Anti-human isotype (IgG3) | AF488 | Biolegend, #401324 | FC | 1:10 |
| Anti-human cTnT (IgG1) | FITC | Miltenyi, #130-119-575 | FC | 1:50 |
| Anti-human isotype control (IgG1) | FITC | Miltenyi, #130-118-354 | FC | 1:50 |

**Table S6:**
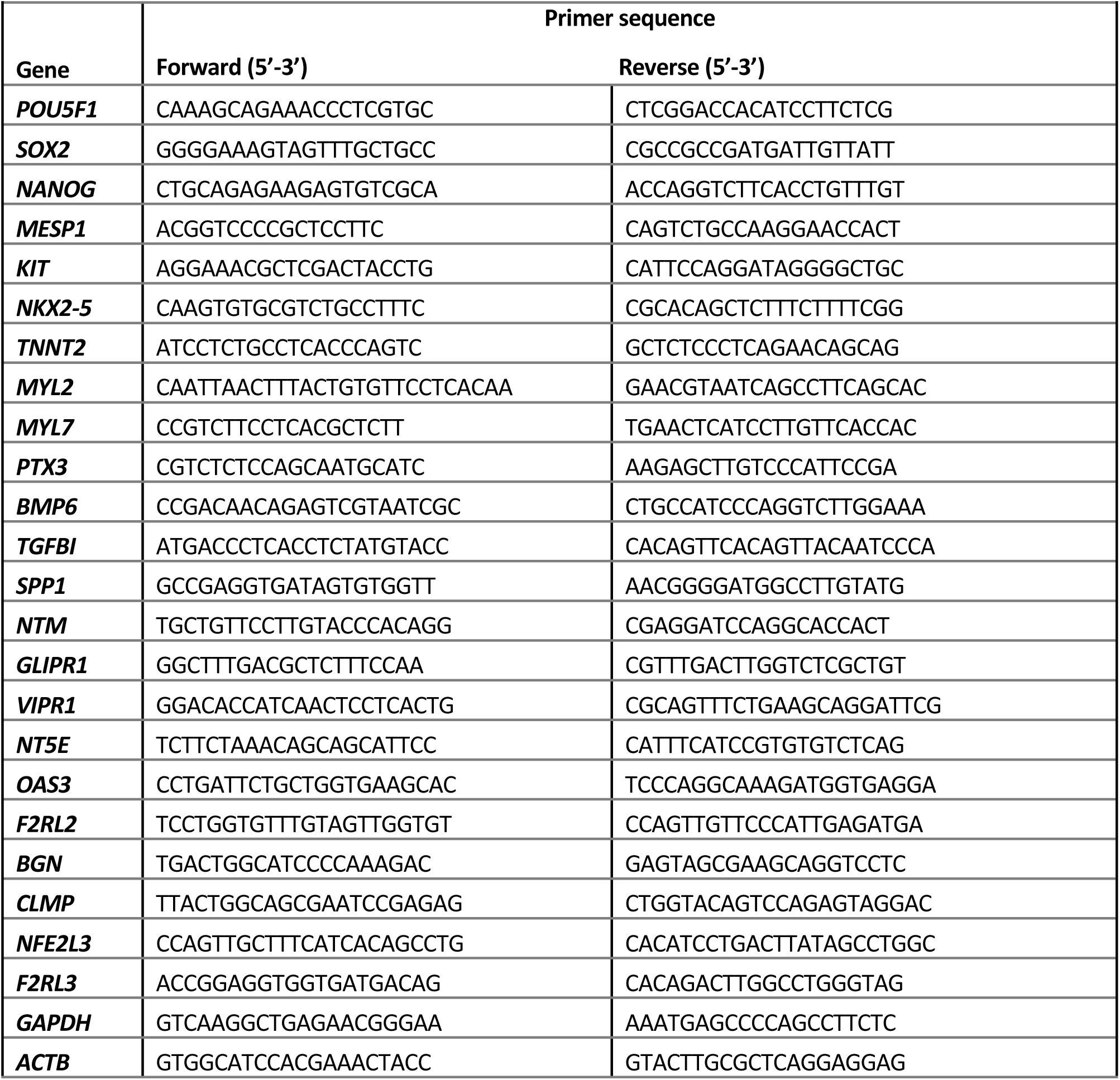
Forward and reverse primer sequences for the genes of interest for RT-qPCR. All primers have a melting temperature of approximately 60°C.

## Bibliography

1. Tsao, C. W. et al. Heart Disease and Stroke Statistics—2023 Update: A Report From the American Heart Association. Circulation 147, E93–E621 (2023).

2. Kormos, R. L. et al. The Society of Thoracic Surgeons Intermacs Database Annual Report: Evolving Indications, Outcomes, and Scientific Partnerships. Ann Thorac Surg 107, 341–353 (2019).

3. Slaughter, M. S. et al. Advanced heart failure treated with continuous-flow left ventricular assist device. New England Journal of Medicine 361, 2241–2251 (2009).

4. Kawamura, T. et al. Safety confirmation of induced pluripotent stem cell-derived cardiomyocyte patch transplantation for ischemic cardiomyopathy: first three case reports. Front Cardiovasc Med 10, 1–9 (2023).

5. Shiba, Y. et al. Allogeneic transplantation of iPS cell-derived cardiomyocytes regenerates primate hearts. Nature 538, 388–391 (2016).

6. Safety and Efficacy of Induced Pluripotent Stem Cell-derived Engineered Human Myocardium as Biological Ventricular Assist Tissue in Terminal Heart Failure. ClinicalTrials.gov identifier: NCT04396899. Updated March 27, 2023. Accessed February 06,2025. https://clinicaltrials.gov/study/NCT04396899

7. Miki, K. et al. Bioengineered Myocardium Derived from Induced Pluripotent Stem Cells Improves Cardiac Function and Attenuates Cardiac Remodeling Following Chronic Myocardial Infarction in Rats. Stem Cells Transl Med 1, 430–437 (2012).

8. Hata, H. et al. Engineering a novel three-dimensional contractile myocardial patch with cell sheets and decellularised matrix. European Journal of Cardio-thoracic Surgery 38, 450–455 (2010).

9. Moyes, K. W. et al. Human Embryonic Stem Cell-Derived Cardiomyocytes Migrate in Response to Gradients of Fibronectin and Wnt5a. Stem Cells Dev 22, 2315–2325 (2013).

10. Wilson, K. D. et al. Endogenous Retrovirus-Derived lncRNA BANCR Promotes Cardiomyocyte Migration in Humans and Non-human Primates. Dev Cell 54, 694–709.e9 (2020).

11. Hosen, M. R. et al. Airn Regulates Igf2bp2 Translation in Cardiomyocytes. Circ Res 122, 1347–1353 (2018).

12. Qin, X., Gao, S., Yang, Y., Wu, L. & Wang, L. microRNA-25 promotes cardiomyocytes proliferation and migration via targeting Bim. J Cell Physiol 234, 22103–22115 (2019).

13. Bergeron, A. et al. Filamentous nestin and nonmuscle myosin IIB are associated with a migratory phenotype in neonatal rat cardiomyocytes. J Cell Physiol 236, 1281–1294 (2021).

14. Bock-Marquette, I., Saxena, A., White, M. D., DiMaio, J. M. & Srivastava, D. Thymosin β4 activates integrin-linked kinase and promotes cardiac cell migration, survival and cardiac repair. Nature 432, 466–472 (2004).

15. Chablais, F. & Jaźwińska, A. The regenerative capacity of the zebrafish heart is dependent on TGFβ signaling. Development 139, 1921–1930 (2012).

16. Peng, Y. et al. Inhibition of TGF-β/Smad3 Signaling Disrupts Cardiomyocyte Cell Cycle Progression and Epithelial–Mesenchymal Transition-Like Response During Ventricle Regeneration. Front Cell Dev Biol 9, 1–14 (2021).

17. Itou, J. et al. Migration of cardiomyocytes is essential for heart regeneration in zebrafish. Development (Cambridge) 139, 4133–4142 (2012).

18. Burridge, P. W., Holmström, A. & Wu, J. C. Chemically Defined Culture and Cardiomyocyte Differentiation of Human Pluripotent Stem Cells. Curr Protoc Hum Genet 87, 21.3.1–21.3.15 (2015).

19. Burridge, P. W. et al. Chemically defined generation of human cardiomyocytes. Nat Methods 11, 855–860 (2014).

20. Lundy, S. D., Zhu, W. Z., Regnier, M. & Laflamme, M. A. Structural and functional maturation of cardiomyocytes derived from human pluripotent stem cells. Stem Cells Dev 22, 1991–2002 (2013).

21. Zuppinger, C. et al. Characterization of cytoskeleton features and maturation status of cultured human iPSC-derived cardiomyocytes. European Journal of Histochemistry 61, 145–153 (2017).

22. Ormrod, B. & Ehler, E. Induced pluripotent stem cell-derived cardiomyocytes—more show than substance? Biophys Rev 15, 1941–1950 (2023).

23. Cappelli, C. et al. The TGF-β profibrotic cascade targets ecto-5′-nucleotidase gene in proximal tubule epithelial cells and is a traceable marker of progressive diabetic kidney disease. Biochim Biophys Acta Mol Basis Dis 1866, 165796 (2020).

24. Dituri, F., Cossu, C., Mancarella, S. & Giannelli, G. The Interactivity between TGFβ and BMP Signaling in Organogenesis, Fibrosis, and Cancer Activated Protein Kinases (MAPKs, namely Extracellular Receptor Kinase 1 and 2-ERK1/2. Cells 8, 1–21 (2019).

25. Hou, J. et al. DACT2 regulates structural and electrical atrial remodeling in atrial fibrillation. J Thorac Dis 12, 2039–2048 (2020).

26. Kramerova, I. et al. Spp1 (osteopontin) promotes TGFβ processing in fibroblasts of dystrophin-deficient muscles through matrix metalloproteinases. Hum Mol Genet 28, 3431 (2019).

27. Lange, A. W., Keiser, A. R., Wells, J. M., Zorn, A. M. & Whitsett, J. A. Sox17 promotes cell cycle progression and inhibits TGF-β/Smad3 signaling to initiate progenitor cell behavior in the respiratory epithelium. PLoS One 4, (2009).

28. Thuault, S. et al. Transforming growth factor-β employs HMGA2 to elicit epithelial-mesenchymal transition. Journal of Cell Biology 174, 175–183 (2006).

29. Li, Q., Xu, Y., Li, X., Guo, Y. & Liu, G. Inhibition of Rho-kinase ameliorates myocardial remodeling and fibrosis in pressure overload and myocardial infarction: Role of TGF-β1-TAK1. Toxicol Lett 211, 91–97 (2012).

30. Vilahur, G. et al. Molecular and cellular mechanisms involved in cardiac remodeling after acute myocardial infarction. J Mol Cell Cardiol 50, 522–533 (2011).

31. Inman, G. J. et al. SB-431542 is a potent and specific inhibitor of transforming growth factor-β superfamily type I activin receptor-like kinase (ALK) receptors ALK4, ALK5, and ALK7. Mol Pharmacol 62, 65–74 (2002).

32. Reibman, J. et al. Transforming growth factor β1, a potent chemoattractant for human neutrophils, bypasses classic signal-transduction pathways. Proc Natl Acad Sci U S A 88, 6805–6809 (1991).

33. Wahl, S. M. et al. Transforming growth factor type β induces monocyte chemotaxis and growth factor production. Proc Natl Acad Sci U S A 84, 5788–5792 (1987).

34. Liu, Z., Yi, L., Du, M., Gong, G. & Zhu, Y. Overexpression of TGF-β enhances the migration and invasive ability of ectopic endometrial cells via ERK/MAPK signaling pathway. Exp Ther Med 4457–4464 (2019) doi:10.3892/etm.2019.7522.

35. Melzer, C., von der Ohe, J., Hass, R. & Ungefroren, H. TGF-β-dependent growth arrest and cell migration in benign and malignant breast epithelial cells are antagonistically controlled by Rac1 and Rac1b. Int J Mol Sci 18, (2017).

36. Zhao, W. et al. Effect of TGF-β1 on the Migration and Recruitment of Mesenchymal Stem Cells after Vascular Balloon Injury: Involvement of Matrix Metalloproteinase-14. Sci Rep 6, 1–11 (2016).

37. Azhar, M. et al. Transforming growth factor beta in cardiovascular development and function. Cytokine Growth Factor Rev 14, 391–407 (2003).

38. Hanna, A. & Frangogiannis, N. G. The Role of the TGF-β Superfamily in Myocardial Infarction. Front Cardiovasc Med 6, 1–15 (2019).

39. Dobaczewski, M., Gonzalez-Quesada, C. & Frangogiannis, N. G. The extracellular matrix as a modulator of the inflammatory and reparative response following myocardial infarction. Journal of Molecular and Cellular Cardiology Preprint at 10.1016/j.yjmcc.2009.07.015 (2010).

40. Yi, X. et al. Hepatocyte growth factor regulates the TGF-β1-induced proliferation, differentiation and secretory function of cardiac fibroblasts. Int J Mol Med 34, 381–390 (2014).

41. Pardali, E. & Dijke, P. Transforming growth factor-beta signalling and tumor angiogenesis. Mol Cell 14, 4848–4861 (2009).

42. Dark, N. et al. Generation of left ventricle-like cardiomyocytes with improved structural, functional, and metabolic maturity from human pluripotent stem cells. Cell Reports Methods 3, 100456 (2023).

